# Implicit adaptation to mirror-reversal is in the correct coordinate system but the wrong direction

**DOI:** 10.1101/2021.05.28.446174

**Authors:** Tianhe Wang, Jordan A. Taylor

## Abstract

Learning in visuomotor adaptation tasks is the result of both explicit and implicit processes. Explicit processes, operationalized as re-aiming an intended movement to a new goal, account for the lion’s share of learning while implicit processes, operationalized as error-dependent learning that gives rise to aftereffects, appear to be highly constrained. The limitations of implicit learning are highlighted in the mirror-reversal task, where implicit corrections act in opposition to performance. This is surprising given the mirror-reversal task has been viewed as emblematic of implicit learning. One potential confound of these studies is that both explicit and implicit processes were allowed to operate concurrently, which may interact, potentially in opposition. Therefore, we sought to further characterized implicit learning in a mirror-reversal task with a clamp design to isolate implicit learning from explicit strategies. We confirmed that implicit adaptation is in the wrong direction for mirror-reversal and operates as if the perturbation were a rotation, and only showed a moderate attenuation after three days of training. This result raised the question of whether implicit adaptation blindly operates as though perturbations were a rotation. In a separate experiment, which directly compared a mirror-reversal and a rotation, we found that implicit adaptation operates in a proper coordinate system for different perturbations: adaptation to a mirror-reversal and rotational perturbation is more consistent with Cartesian and polar coordinate systems, respectively. It remains an open question why implicit process would be flexible to the coordinate system of a perturbation but continue to be directed inappropriately.

**Public Significance Statement:** Patients with severe amnesia can improve their performance from day to day in mirror-reversal tasks. These findings led, in part, to the codification of explicit and implicit processes in classic theories regarding the taxonomy of memory systems, with motor learning resting firmly in the branch of implicit memory. However, recent evidence has shown that explicit processes also play an important role in motor learning. What’s more, these studies have found that implicit learning doesn’t operate in a useful way in the mirror-reversal task. In the present *study*, we further examine this puzzling behavior of implicit learning in a mirror-reversal task using a design that can isolate implicit processes from explicit strategies. We clearly showed that the implicit system adapts in the wrong direction for a mirror-reversal, acting as if the perturbation were a rotation. Surprisingly, however, we found that although adaptation is in the wrong direction, the implicit system is sensitive to a particular coordinate system. These findings further challenge the flexibility of this implicit adaptation process in motor learning.

## Introduction

Since the seminal findings of Patient H.M., there has been a tendency to view motor learning as the result of an implicit process (Milner, 1962; Corkin, 1968; Squire, 2004). However, in recent years, it has become clear that motor learning can result from a number of different learning processes (Heuer & Hegele, 2008, 2010; Izawa & Shadmehr, 2011; Verstynen & Sabes, 2011; Taylor, Krakauer, & Ivry, 2014). As such, there has been considerable interest in isolating implicit learning from these other processes to better characterize its operation (Mazzoni & Krakauer, 2006; Taylor & Ivry, 2011; Morehead et al., 2017; Kim et al., 2018; Leow, Marinovic, de Rugy, & Carroll, 2018). One of the most surprising findings from these efforts to study implicit learning, in the context of a visuomotor adaptation task, is that it appears to have a very limited dynamic range: It is relatively insensitive to the magnitude of the perturbation (Bond & Taylor, 2015; Morehead et al., 2017; Hutter & Taylor, 2018), only corrects for errors in one direction (Butcher & Taylor, 2018), operates automatically regardless of task goals (Morehead et al., 2017; Kim et al., 2018), saturates at a value far below complete learning (Bond & Taylor, 2015; Wilterson & Taylor, 2019), and may even reduce its contribution with extended training (Avraham, Morehead, Kim, & Ivry, 2021).

These findings present a significant challenge to current theories of sensorimotor adaptation, which center around the idea of an internal forward model (Jordan & Rumelhart, 1992; Krakauer & Shadmehr, 2006; Shadmehr, Smith, & Krakauer, 2010; Hadjiosif, Krakauer, & Haith, 2021). Indeed, it has been suggested that implicit adaptation process observed in visuomotor adaptation tasks may not reflect the operation of an internal forward model and, perhaps, it should be considered the outcome of a different computational process (Wolpert, Ghahramani, & Flanagan, 2001; Abdelghani, Lillicrap, & Tweed, 2008; Krakauer et al., 2019; Wilterson & Taylor, 2019; Avraham et al., 2021; Hadjiosif et al., 2021). In fact, it is the mirror-reversal task used with Patient H.M. that underscores the limitations of this implicit process. Upon experiencing a mirror reversal, feedback corrections appear to be in the wrong direction and this response only gently decreases with extended training (Gritsenko, Yakovenko, & Kalaska, 2009; Telgen, Parvin, & Diedrichsen, 2014; Kasuga et al., 2015). Likewise, the adaptive response is similarly in the wrong direction, creating trial-by-trial instabilities in performance (Hadjiosif et al., 2021), and only becomes suppressed with days of training (Wilterson & Taylor, 2019). Thus, it appears that, at least initially, implicit motor adaptation shows a highly stereotyped response that is incompatible with the operation of a forward model (Hadjiosif et al., 2021).

While it might be tempting to reach this conclusion based on the recent findings in the mirror-reversal task, it should be noted that in these previous studies learning may be contaminated by other processes, such as explicit re-aiming strategies. Recent studies have found that explicit strategies warp the generalization of implicit learning (Day et al., 2016; McDougle et al., 2017), task performance errors interact with implicit adaptation processes (Kim et al., 2018; Leow et al., 2018), and how explicit and implicit processes are measured may influence the observed pattern of results (Maresch, Werner, & Donchin, 2021). While it seems unlikely that the operation of other learning processes, or the way in which they are assayed, could dramatically alter the underlying operation of implicit adaptation process, such as reversing the direction of corrective or adaptive response, it still remains worthwhile to attempt to isolate it.

The task-irrelevant visual-error clamp task has recently been developed with the hopes of isolating implicit adaptation from other sources of potential contamination (Morehead et al., 2017). Here, in a center-out visuomotor-rotation task, the angular direction of cursor feedback is fixed regardless of hand direction. Participants have control only over the radial direction of the movement and are instructed to ignore cursor feedback entirely. Despite the task-irrelevance of the feedback, movements gradually begin drifting in the direction opposite of the cursor -- most often without awareness of the participants.

While these initial studies applied a rotational perturbation to the cursor, the perturbation can equally arise from a mirror reversal, thus, presenting the opportunity to study implicit adaptation in a mirror reversal task in relative isolation from other processes. Here, we employed this mirror-reversal clamped-feedback task to better characterize implicit adaptation over the course of three consecutive days of training. Regardless of the participants’ reach direction, cursor feedback was always reflected across the mid-sagittal axis of the workspace (i.e., x-axis). Participants were informed that the movement of the cursor was not under their control, so they should ignore it and try their best to place their hand under the presented target. Similar to previous reports of implicit adaptation under a mirror reversal, we found that implicit adaptation was in the inappropriate direction and only gently decreased across sessions. What’s more, the observed pattern of adaptation was consistent with what we would be expected of adapting to different rotation magnitudes and directions across the workspace, in effect, reflecting a complex in a dual-adaptation paradigm to opposing rotations (Schween & Hegele, 2017; Schween, Taylor, & Hegele, 2018).

The finding that adaptation to a mirror reversal is homologous to a rotation raises the question: Does implicit adaptation truly operate as though visual errors arise from a rotational perturbation? That is, does adaptation operate in a polar-like coordinate system? This question has been asked before by Hudson and Landy (2016) in a paradigm where the errors were either drawn from a polar or Cartesian-based coordinate system. While they found that the adaptive response was flexible to either form of perturbations, they did not make a dissociation between explicit or implicit learning processes. Thus, we sought to revisit this question contrasting adaptive responses to mirror-reversal and rotational perturbation using a paradigm that assays both implicit and explicit learning (Bond & Taylor, 2017; Hutter & Taylor, 2018). While we again found that the direction of implicit adaptation to a mirror reversal was similar to a rotation, the pattern of the adaptive response appeared to be appropriate for the coordinate system.

## Experiment 1 (Clamped mirror-task)

### Materials and Methods

#### Participants

Fourteen young adults (six women, age: 20.5 ± 0.5y) were recruited from the research participation pool maintained by the Department of Psychology at Princeton University or from the local community. One participant withdrew midway through the study, resulting in thirteen participants for analysis. All participants were right-handed, verified by the Edinburgh handedness test (Oldfield, 1971) and reported normal or corrected-to-normal visual acuity. The experiment consisted of separate 1-hour sessions spread across three days of participation. Participants performed each session approximately at the same time each day. All participants provided informed, written consent before participation. The study protocol was reviewed and approved by the Princeton University’s Institutional Review Board. All participants received $60 for participation.

#### Apparatus

Participants performed a center-out reaching task, making horizontal movements with their right hand. Movements were recorded by a digitizing tablet (Wacom Co., Kazo, Japan), which recorded the motion of a digitizing pen held in the hand. Stimuli were displayed on a 60 Hz, 17-in. touch-sensitive monitor (Planar Systems, Hillsboro, Oregon), which was mounted horizontally above the tablet and, thus, obscured view of the limb. The task was controlled by custom software coded in MATLAB (The MathWorks, Natick, Massachusetts), using Psychtoolbox extensions (Brainard, 1997, Kleiner, Brainard, and Pelli, 2007), and run on a Dell OptiPlex 7040 computer (Dell, Round Rock, Texas) with Windows 7 operating system (Microsoft Co., Redmond, Washington).

#### Procedure

Each trial began with participants holding their right hand in the center of the workspace for 500 ms. After this delay, a green target (0.25 cm radius) appeared 7 cm from the start location. The target could appear at one of twelve locations (+/−30°, +/−60°, +/− 75°, +/−105°, +/−120°, +/−150°; see Figure 1A). Participants were instructed to move as quickly as possible to the target and then immediately return back to the center in one smooth movement. During the outbound portion of the reach, a small circular cursor (0.15 cm radius) was displayed. Depending on the phase of the experiment, the cursor either represented the location of the hand or clamped feedback (see below). In addition, a neutral “click” sounded to indicate that the participant’s hand exceeded 7 cm. If the movement time from the start to 7 cm exceeded 800 ms, then a computer-generated voice sounded “Too Slow” following the “click” (<1% of trials, which were excluded from further analysis). During the inbound portion of the reach, the small circular cursor was replaced by a white ring that was centered on the start location and with a radius that matched the distance of the participants’ hand to the start location. This form of feedback was used to help guide participants back to the start location by providing only radial but not directional information.

**Figure 1.**
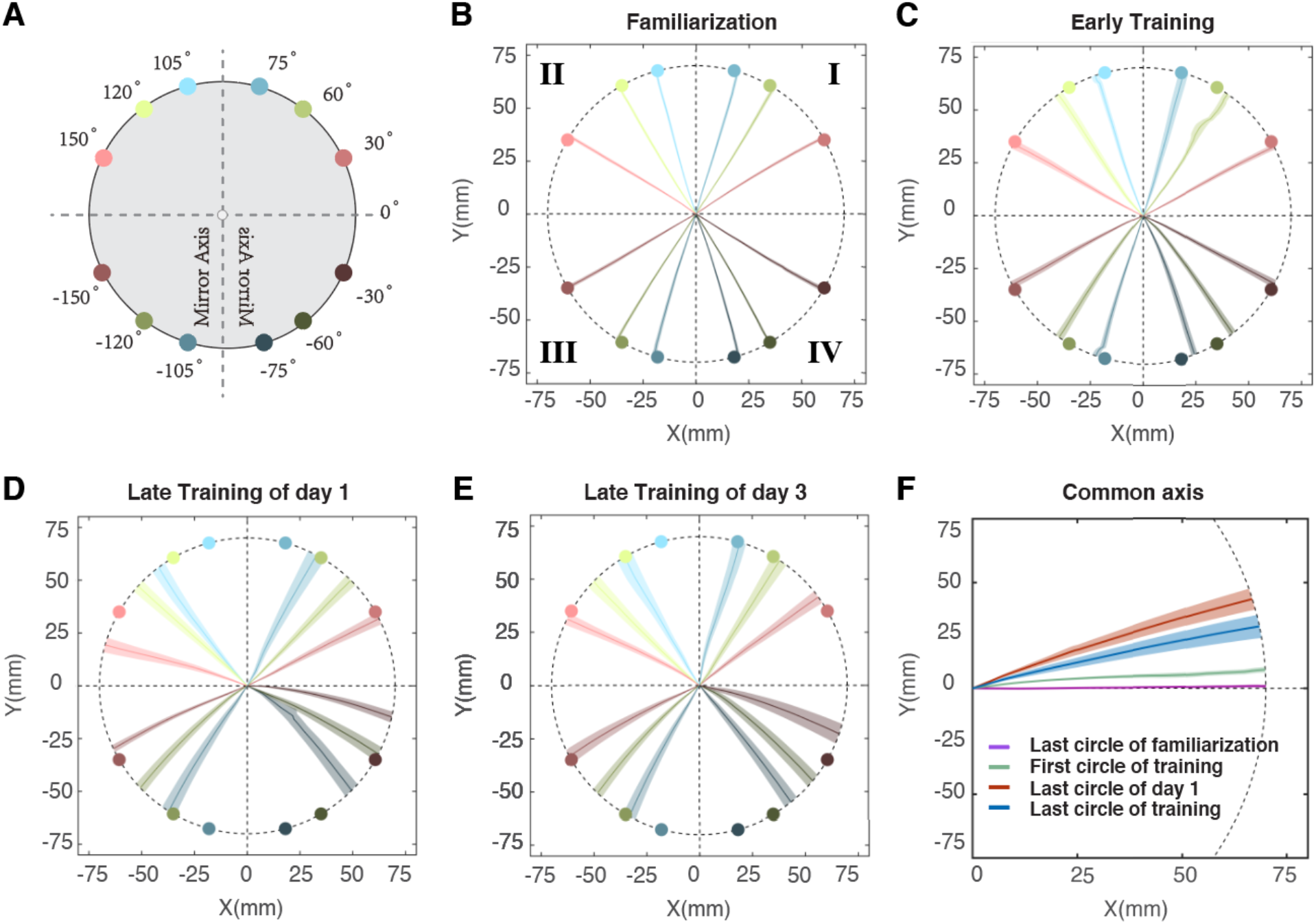
Target locations and moving trajectories of Experiment 1. A) The locations of twelve targets: +/−30°, +/−60°, +/− 75°, +/−105°, +/−120°, +/−150°. Mirror-reversal was applied along the y-axis and centered on the starting location. Targets are color-coded by the size of error between clamped feedback and target location: red: 120°, green: 60°, blue: 30°. Different darkness levels indicate 4 quadrants. B) Moving trajectory of the last circle in the familiarization phase. Each trajectory is averaged over thirteen participants and colored according to the target it aims. The shaded area represents standard error. Same below. C) Moving trajectory of the first circle in the clamped phase of day one. D) Moving trajectory of the last circle in the clamped phase of day one. E) Moving trajectory of the last circle in the clamped phase of day three. F) Averaged Moving trajectories of B-E. All twelve targets in one circle are flipped and rotated into a common axis at 0° (see methods) and averaged.

Participants received either veridical feedback or clamped feedback in different phases of Experiment 1. On veridical feedback trials, the cursor accurately showed the location of the hand. For visual-error clamp feedback trials, the feedback followed a trajectory that was fixed along a specific heading angle (Shmuelof, Krakauer, & Mazzoni, 2012; Morehead et al., 2017). This fixed heading angle was determined as the mirror-reversed direction of the target angle across the vertical axis (mid-sagittal axis). The radial location of the cursor was based on the radial extent of the participant’s hand (up to 7 cm amplitude) but was independent of the angular location of the hand.

On the first day of the experiment, participants experienced a familiarization phase consisting of 60 trials with veridical, online feedback. Following the familiarization phase, the experimenter then paused the task to explain the visual-error clamped feedback. Participants were informed that they could no longer control the direction of the cursor movement in the following sessions of the experiment. They were instructed to ignore the cursor and continue to reach directly to the visual location of the target. Participants then practiced completing twelve trials, one to each target location, with the clamped feedback. The experimenter then paused the experiment again to ensure that the participants understood the instructions and confirmed that they understood that they could not – and should not – attempt to control the direction of the cursor. Following this practice phase, participants completed another 996 trials with clamped feedback (clamp phase). On the second day of the experiment, participants completed another 1008 trials of clamped feedback, which was present on the very first trial. On the third day of the experiment, participants completed 1500 trials of clamped feedback and ended the experiment with a “washout” of 60 trials without any feedback. In sum, participants completed 3636 trials in total and note, they were instructed to aim at the visual target directly in all phases of the experiment.

Data analysis — All initial data analyses were conducted in Matlab, with the exception that repeated measures ANOVAs were conducted in SPSS (IBM, 2011). The Greenhouse-Geisser correction was used for ANOVAs when data violated the homoscedasticity assumptions. Multiple comparisons were Bonferroni corrected. Task performance was assessed by calculating the angular difference between the target and the initial heading angle of the hand. The initial heading angle of the hand was calculated as the angle between the first and last sample of hand position between 1 and 3cm radially from the start location.

Our previous study employing a mirror-reversal found that the direction of the implicit component of motor adaptation was not in a direction consistent with counteracting a mirror-reversal but instead in a direction consistent with countering a rotation (Wilterson & Taylor, 2019). As such, this requires a relatively unintuitive transformation of the data to get it to a common axis. Figure 1D presents the raw hand trajectory of the last ten trials toward each target at the end of the first-day training. It is clear from this figure that to transform the data to a common axis, the movements in quadrants II-VI need to be transformed as follows: II) the x-locations of the second quadrant movements (targets 105°, 120°, 150°) were multiplied by −1 to flip the y-axis, III) both x- and y-locations of the third quadrant movements (targets −105°, −120°, −150°) were multiplied by −1, and IV) the y-locations of the fourth quadrant movements (targets −30°, −60°, −75°) were multiplied by −1. After this, the flipped trajectories were rotated to a common axis, as though the intended target was always located at 0°, despite being oriented at different target directions. With these transformations, a positive angle indicates movements of the hand in a counterclockwise direction relative to the target.

The order of the targets was pseudorandomized, such that each of the twelve targets was visited once before being repeated. To minimize the influence of target order differences between subjects and the potential biomechanical biases associated with specific target directions, trials were averaged into bins of twelve trials. In addition, the averaged hand angle of the last three bins (36 trials) in the clamp phases was calculated for day 1-3, respectively, to assess the adaptation of training of each day. The hand angles in the clamp phase and washout were corrected by subtracting with the mean of the familiarization phase.

#### Power Analysis

In our previous five-day mirror-reversal task (Wilterson & Taylor, 2019), the magnitude of implicit adaptation reached the peak at the end of the first day and gradually decreased over the course of the next four days of training. Based on these findings, it is reasonable to hypothesize that the adaptation in the early phase of the training under the clamped mirror also decreases after multiple days of training. To decide the necessary sample size we need to examine this hypothesis, the means of implicit adaptation at the end of day one were compared with the end of day 2, day 3, day 4, and day 5 in our previous five-day mirror task (32 trials based on Wilterson & Taylor, 2019). Effect sizes of d = 0.99, 1.08, 1.08, 1.53 were found for day 2, day 3, day 4, day 5 compared to day 1, respectively. A paired t-test using a two-tailed α of 0.05 and power of 0.8 suggested that a sample size of 17, 14, 14, 7 participants was needed, respectively. Considering the constraints in participant recruitment, we decided to perform a three-day experiment with 14 participants, which was chosen as a balanced compromise in terms of length of the experiment, power, and typical sample sizes is in visuomotor adaptation experiments.

## Results and Discussion

Previous studies of adaptation under mirror-reversed feedback have found that feedback corrections and adaptation are in a direction that is counterproductive to overcoming the mirror perturbation. Instead, the responses appear to be more consistent with an adaptation system that treats the perturbation as though it was a rotation. However, as training progresses the inappropriate response is gradually decreased. Here, we set out to study this behavior in more detail by isolating implicit adaptation over the course of three days of training under clamped mirror feedback.

### The direction of adaptation

Prior to the introduction of the clamped mirror feedback, participants practiced reaching to the targets with veridical feedback. Their reaches were fairly accurate in reaching toward the target (0.94 ± 0.74°, mean ± standard deviation, same below, Figure 1B). Following this familiarization phase, cursor feedback was mirror-reversed and clamped, where the mirror-reversal was applied along the y-axis and centered on the starting location and the angle of the cursor feedback was fixed.

Initially, participants reaching movements were directed toward the target, but with training their reaches deviated from the target (Figure 1C/D). For targets in all four quadrants, the heading angle of reaching movements biased considerably away from the mirror-axis at the end of the first day (Figure 1D) and this bias was still present at the end of the third day of training (Figure 1E). Specifically, the heading angle drifted clockwise in quadrant I and quadrant III, and drifted counterclockwise in quadrant II and quadrant IV. When considering the target location in the workspace relative to mirror axis, this pattern of results suggests that implicit adaptation is acting to in a direction more consistent with adaptation to a visuomotor rotation rather than a mirror reversal as we have observed previously (Wilterson & Taylor, 2019; Hadjiosif et al., 2021). Indeed, when we transformed the data to a common axis (see Methods), it is clear that implicit adaptation is acting as if to counter a visuomotor rotation (Figure 1F).

Figure 1F also reveals that the amount of adaptation on the final day of training is less than that observed at the end of the first day. With the data now on a common axis, it is evident that the amount of implicit adaptation peaked at the end of the first day (15.8 ± 6.3°) and only gently decreased over the remaining two days of training (Figure 2A); by the end of day two, adaptation was 13.6 ± 6.9° and by the end of day three was 10.8 ± 7.1°. This decrease was confirmed by submitting these data to a three-way, repeated measures ANOVA with the factor of Day ANOVA: F_2, 24_ = 8.32, p = 0.002, 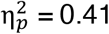 (Figure 2B). While there is a significant decrease over the course of three days, the decrease is quite modest averaging only 2-3° each day. In the post hoc comparisons, only day 1 and day 3 showed a significant difference (p=0.02). At this pace, it would take an additional four days of training to decline to zero – a total of seven days of training with approximately 7000 trials. However, it appears as though the reduction occurs between days rather than within a day. To test this possibility, we performed a separate linear regression on the time series of hand angles for each day of training to determine if there was a significant slope to the time course. As expected, we found a positive slope in day 1 (0.077 ± 0.068 °/trial, t_12_ = 4.08, p = 0.002), and almost a zero slope in day 2 (0.0007 ± 0.0406 °/trial, t_12_ = 0.06, p = 0.95) and day 3 (−0.007 ± 0.027 °/trial, t_12_ = −0.90, p = 0.39), indicating almost no decrease within a day.

**Figure 2.**
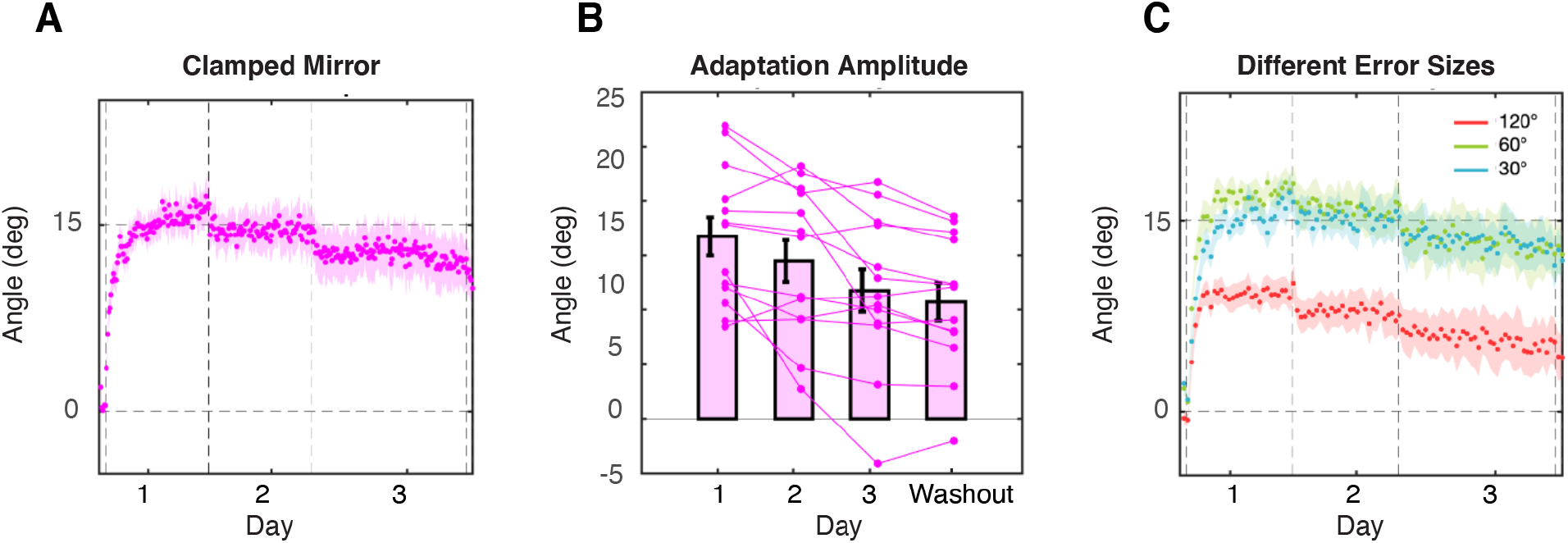
Time course of the adaptation in Experiment 1. A) Time course of the hand angle over twelve targets across three days. Each dot is an average of twelve trials (one circle). The horizontal dashed lines represent first the onset of the mirror-clamp perturbation, then day breaks, then the start of washout. The shaded area represents standard error. Same below. B) Amplitude of adaptation after each day training. The error bars indicate mean and standard error. C) Time courses of the hand angle for three different error sizes. Each dot is the average of twelve trials (three circles).

While we observed a reduction in the amount of implicit adaptation over the course of three days of training, it remains an open question as to whether the adaptive response of the implicit system would become appropriate for a mirror reversal or simply be suppressed. Note, if the adaptation was in the mirror direction, i.e., the adaptation worked to bring the cursor to the target following the rule of a mirror, the hand would have drifted to the direction of the cursor. The cursor was in the counterclockwise direction to the targets in quadrant I (the clamped feedback for them was in quadrant II), so the hand angle would be negative instead of positive. Nevertheless, after the three days of training, only one participant showed a slight negative hand angle at the end of day three, while all the other participants’ hand angles remained positive throughout the whole training process (Figure 2B).

In the washout phase with no visual feedback, implicit adaptation was still present (9.8 ± 6.2°), which was slightly smaller than the end of the training session (t_12_ = 2.46, p = 0.03). Notably, before the washout phase, we needed to pause the program to give participant instruction, which resulted in a longer intertrial interval (11.3 ± 3.8s) than the normal intertrial interval in the training phase (1.2 ± 0.1s). This may be one reason why adaptation in washout is smaller than at the end of training.

### Influence of error size

The finding that implicit adaptation under a mirror reversal resembles that observed under a visuomotor rotation and that previous studies have revealed that the extent of generalization of a rotation is narrow, then we should expect that each target direction is learned relatively independently. Furthermore, if the mirror-reversal task is acted upon by the implicit system as a rotational perturbation, then each target would technically have a different degree of rotation. For example, reaches to the 30° target (quadrant I) results in errors near the 150° target (quadrant II), which could be viewed as a 120° rotation. Likewise, reaches to the 75° target (quadrant I), results in errors at 105° (quadrant II), which could be viewed as a 30° rotation. Morehead and colleagues (2017) found that in a rotation-based visual-error clamp, adaptation declines as the size of the rotation increases (adaptation decreased at a point between 95° and 135°). Thus, if implicit adaptation treats mirror reversal perturbations as a rotational error, then we would expect more adaptation for targets resulting in smaller error (<95-135°). Indeed, when we separate the learning curves by targets resulting in similar errors, the adaptation for different error sizes showed a significant difference after the first day of training (F_2, 24_ = 27.2, p < 0.001,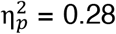). Post hoc pairwise comparisons showed that the adaptation to target of 120° errors (targets +/−30, +/−150; 9.9 ± 3.3°) was significantly smaller than the adaptation to targets of 60° errors (targets +/−60, +/−120; 16.3 ± 4.9°, p < 0.001) or the adaptation to targets of 30° errors (targets +/−60, +/−120; 15.6 ± 5.4°, p = 0.002). No difference was found between the adaption to 60° errors and the adaptation to 30° errors (p = 0.504), consistent with Morehead and colleagues (2017).

The findings from Experiment 1 suggest that implicit adaptation, even under a mirror reversal, acts as though the error arises from a rotational perturbation: The implicit system adapted in a direction opposite to the visual error and the magnitude of adaptation was target-dependent, which is consistent with a rotational-based error signal. This raises the question: Does the implicit system truly operate in a rotational manner, such as being transformed into a polar-like coordinate system, or does it simply act to bring visual feedback toward the intended reach direction, such as an equal and opposite corrective behavior, which would be more similar to a Cartesian-based system. Note, we do not necessarily think that adaptation actually operates in such a well-defined coordinate system but may be more consistent with a particular coordinate system. In fact, Hudson and Landy (2016) asked this question in a clever visuomotor adaptation task that sought to dissociate these two coordinate systems. Interestingly, they found that participants appropriately adapted to perturbations generated from either coordinate system.

At face value, our current results appear to be at odds with the findings from Hudson and Landy (2016) since implicit adaptation appeared to treat the mirror reversal as a rotation, but their study did not distinguish between explicit and implicit contributions. Explicit re-aiming strategies have been shown to be quite flexible to the details of the perturbation (Bond & Taylor, 2015; Bond & Taylor, 2017; Butcher & Taylor, 2018). While the nature of their perturbations would seem unlikely to afford a large contribution from explicit re-aiming, it still remains an open question. Experiment 1 was ill-equipped to shed light on this issue because it used a center-out task design with shooting movements that always passed the target location (7cm), which prevents the potential coordinate system underlying adaptation to be observed.

Here, we sought to revisit this question by using a modified version of the visual-error clamp method in which participants were required to stop on the target without visual feedback. Once the movement terminated, clamped mirror-reversal endpoint feedback was provided. To dissociate explicit and implicit learning, participants were asked to report their intended reaching location (re-aiming) using a touch-screen monitor before starting the movement. As such, this method would allow us to observe both the x- and y-positions of the hand to observe if learning was more consistent with a rotational correction (e.g., more Polar-like coordinates; Figure 3A) or with an equal and opposite correction (e.g., more Cartesian-like coordinates; Figure 3B).

**Figure 3.**
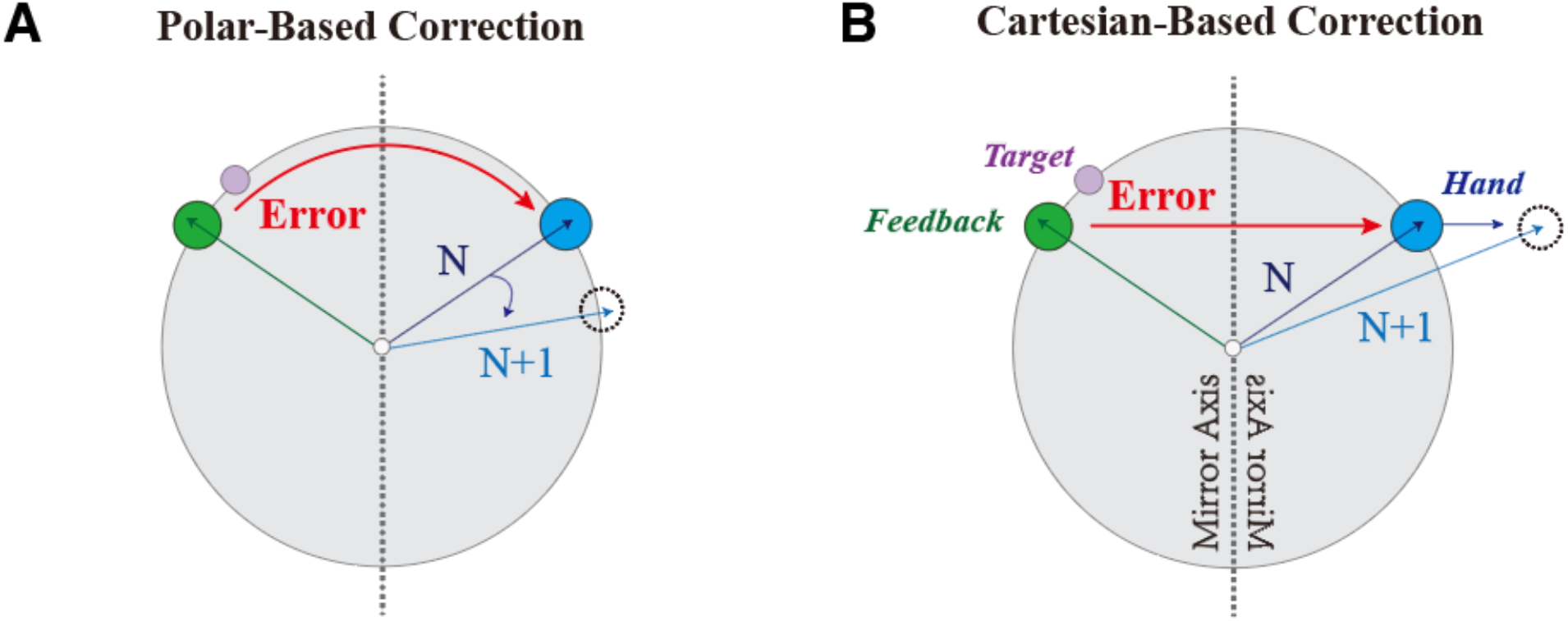
Illustration of correction and target locations in Experiment 2. Appropriate correction of implicit adaptation for Polar (A) and Cartesian (B) coordinate.

## Experiment 2 (Mirror verse Rotation)

### Materials and Methods

#### Participants

Twenty-four participants (thirteen women, age: 23.2 ± 4.3y) were randomly assigned into two perturbation conditions (mirror and rotation), with twelve participants in each condition. Since one participant failed to understand the instruction in the washout phase and continuously aimed at the solution instead of the target, she was excluded from the analysis of aftereffects. Each participant received $12 for participation.

#### Procedure

Each trial began with participants holding their right hand in the center of the workspace for 500 ms. After this delay, a circular orange target (0.25 cm radius) appeared 7 cm from the start location. The target could appear at one of four locations (45°, 135°, −135°, and −45°). The order of the targets was pseudorandomized, such that each of the four targets was visited once before being repeated. To begin the trial, participants indicated their intended reaching position by tapping on the screen with their left hand (Bond & Taylor, 2017; Hutter & Taylor, 2018). Once an aim was recorded, the target turned from orange to green, and participants were allowed to begin their reaching movement with the right hand. If a participant attempted to begin moving their right hand before an aiming location was registered, the message “Remember to report aim” was displayed and the trial restarted. Participants were instructed to move quickly and, different from Experiment 1, they are instructed to stop either on the target or the aim position depended on task phases. If the location of the cursor overlapped with the target (< 2 mm of deviation), the participant heard a pleasant “ding”. Otherwise, an unpleasant “buzz” sounded. The feedback “Too Slow” was given if the reaching time exceeded 1200 ms. After participants stopped moving their right hand (speed < 1.3mm/s), a small circular cursor (0.15-cm radius) appeared to provide feedback for 500 ms. During the perturbation phase, either a mirror-reversal or a visual rotation was applied. In mirror-reversal, the endpoint feedback was reversed over the y-axis. In rotation, the endpoint feedback was rotated 90° to the clockwise direction to match the size of the perturbation in the mirror-reversal task. To avoid priming participants with a certain coordinate system, no visual guidance was used to help participants get back to the center of the workspace. Instead, a small sticker was attached to the tablet surface to provide a tactile cue of the start location. The next trial began once the subject reached the start location.

Sixteen trials with online feedback began the experiment. Then, an 80-trial familiarization phase with veridical endpoint feedback helped participants learn the distance between targets and start location. After that, a baseline phase was composed of 32 trials without feedback to determine if any subject held strong biomechanical biases that would not be averaged out by the target set. Participants were not asked to report their aim during the familiarization phases and baseline phase, as it was assumed that they would always be aiming at the target. A pause was included after the baseline phase so that the experimenter could explain the aiming procedure to the participants. Following these instructions, participants completed 32 trials with veridical feedback to familiarize themselves with the touchscreen and aiming procedure. A perturbation phase of 400 trials with either mirror-reversal or visual-rotation feedback followed this familiarization. The experiment ended with a no-feedback and no-aiming-report washout phase with 32 trials.

#### Data analysis

During the experiments, the digitizing tablet logged the trajectory of the right hand and the touchscreen monitor recorded the location tapped to indicate aiming location. The first data point collected after the movement’s speed was lower than 1.3mm/s was registered as the hand location. Aiming location for each trial was the location tapped on the touchscreen.

In the mirror task, all the hand and aiming locations were transformed such that the locations of the targets were in quadrant II (see method of Experiment 1). Similarly, to make the solutions to perturbations positive, in the rotation task, hand and aiming locations of targets in I, II, and VI were rotated for 180°, 90°, and 270° clockwise, respectively, to match the target in quadrant III. Trials were averaged into bins of 8 trials for each individual (2 cycles). The last 36 trials were used to measure participants’ performance at the end of the perturbation phase. The hand angles in the perturbation phase and washout were corrected by subtracting with the mean of the baseline phase.

Model Simulation — To predicts the time courses of the Cartesian-Based-Correction Model and the Polar-Based-Correction Model, we used a modified version of the two-state model (Smith et al., 2006). Here, we modeled explicit re-aiming as the fast process (Xf) and implicit adaptation as the slow process (Xs) over the course of 624 trials (McDougle et al., 2015). Besides, we assumed that explicit re-aiming is updated based on target error, while implicit adaptation is updated based on the aim-to-cursor distance (McDougle et al., 2015; Equations 1 and 2).

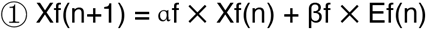

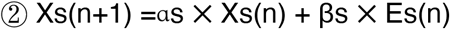

Where Ef is the error between the target and the cursor locations, while Es is the error between explicit re-aiming (Xf) and the cursor location. In the Polar-Based-Correction Model, the errors (Ef, Es) are supposed to be coded in theta-dimension, thus the correction (Xf, Xs) is also performed in theta-dimension. Everything remained zero in the rho-dimension. In the Cartesian Based-Correction Model, Xf, Xs, Ef, and Es were all coded in x-dimension and everything remained zero in the y-dimension. Since our major interest focused on implicit adaptation during the training, rather than how the participants solved the perturbation with explicit strategies, the models assumed that the explicit system can solve the mirror task as in Wilterson & Taylor, (2019). That is, the participants could figure out the solution to explicit re-aiming (Xf) after observing perturbation for one trial. In the following trials, explicit re-aiming (Xf) was updated according to the equations mentioned above. The values for αf, βf, αs, and βs were determined by hand-tuning these parameters to the simple rotation case (αf=1, βf=0.0005, αs=0.99, and βs=0.003). The same parameters were used for the two models. Note, both models assumed the slow process (Xs) updates to compensate mirror-reversal as if it were visuomotor rotation. The simulated data were processed with the same procedure as the behavioral data to make the figures comparable. Note, for simplicity, we opted to present both the model simulations and the behavioral analyses in the Cartesian coordinate system (instead of polar) because if learning in the mirror reversal task is sensitive to the coordinate system, then the majority of learning would be confined to the x-dimension (Figure. 4).

**Figure. 4.**
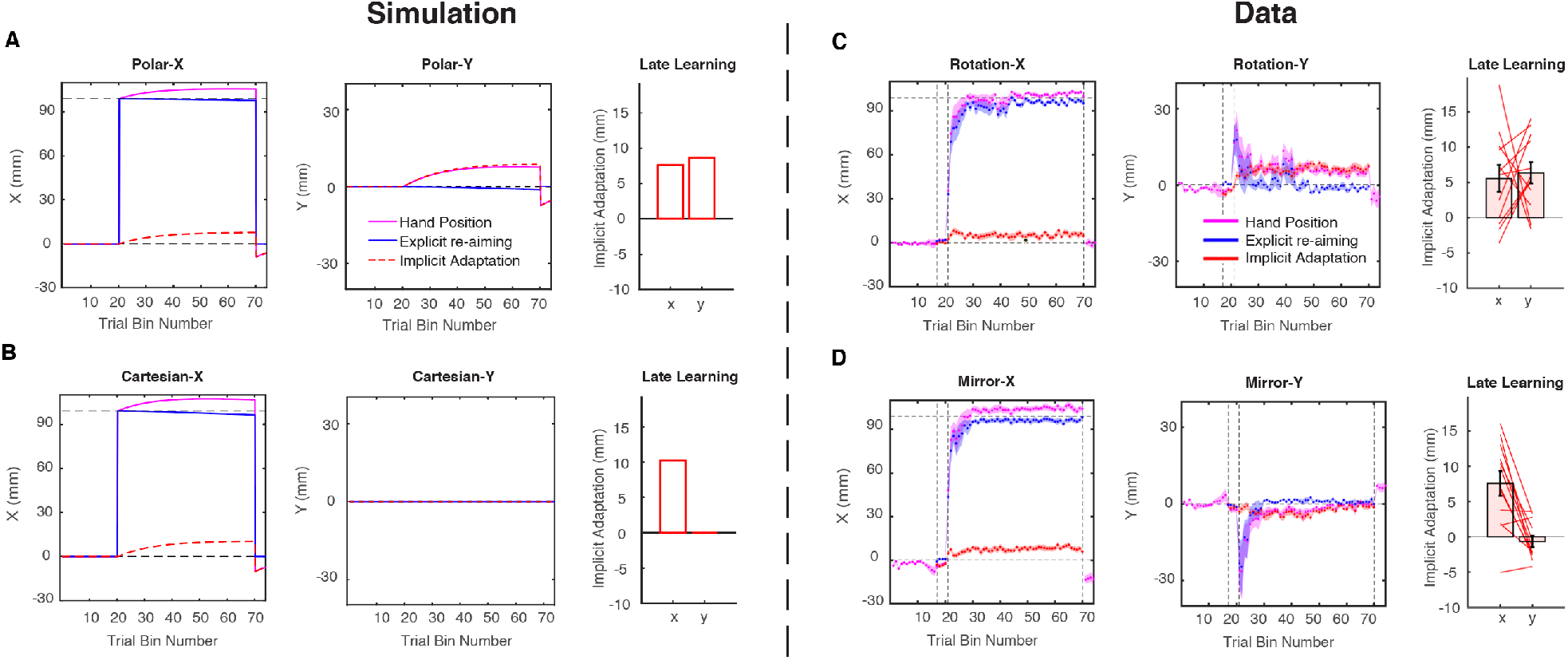
Time courses of model predictions and behavioral result in Experiment 2. (A-B) Simulations of the Polar-based-correction model (A) and the Cartesian-based-correction model (B). Left and middle: Time courses presented by the changes in x- and y-dimensions, respectively. Right: The x- and y-values of implicit adaptation at the end of the perturbation phase. (C-D) Participant’s performance in the rotation task (C) and the mirror task (D). Left and middle: The time course for hand location (purple), cognitive strategy/explicit re-aiming (blue), and implicit adaptation (red). The vertical dashed lines represent the onset of aiming report, then the onset of the visual perturbation, then the end of visual perturbation. The horizontal dashed lines represent the value of 0 mm and 99.0 mm, respectively, which were the perfect solutions of visual perturbation in y- and x-dimensions, respectively. The shaded areas represent standard error. Right: The x- and y-values of implicit adaptation at the end of the perturbation phase. Each line represents a participant. Error bar represents standard error.

#### Power Analysis

Our goal here was to determine how many participants would be needed to detect significant adaptation following a single day of training. We based this estimate on our previous study (Wilterson & Taylor 2019), which found an effect size of d=1.4 for the mean of implicit adaptation at the end of one day of training. A paired t-test using a two-tailed α of 0.05 and power of 0.8 suggested that a sample size of only 7 participants was needed. Since this estimate appeared small, we decided to recruit 12 participants for each task, which is a typical sample size is in visuomotor adaptation experiments and matched that used in our previous study.

## Results and Discussion

### The coordinate system of adaptation

Both groups of participants were able to hit the target precisely in the familiarization phase (Rotation: X −0.7 ± 3.7 mm, Y −1.4 ± 2.7 mm; Mirror: X −1.9 ± 2.7 mm, Y −0.3 ± 1.9 mm). Although some weak biases were generated in the x-axis during the no-feedback baseline phase in the mirror perturbation group (−5.8 ± 8.2 mm), this bias was quickly washed out when the veridical endpoint feedback came back at the beginning of the aiming report (−1.9 ± 5.0 mm). After the participants became familiar with the aiming-report procedure, the perturbation was introduced (mirror-reversal or 90° rotation). In both tasks, participants appeared to solve the perturbations relatively quickly. The hand angle is close to the perfect solution for each perturbation at the end of training (Table 1 – row one). In both tasks, this compensation was carried out in a quite explicit way, since participants could aim precisely at the perfect solutions to the perturbations without any systematic bias at the group level (Table 1 – row two). Although explicit strategy itself is almost sufficient to meet with the task command, the hand positions still deviated away from the aiming positions, indicating the involvement of implicit adaptation (Table 1 –row three).

**Table 1.** The hand location, aiming location and implicit adaptation at the end of the mirror-reversal or rotation perturbation in Experiment 2 (mean (SD); mm).

|  | Rotation-X | Rotation -Y | Mirror -X | Mirror -Y |
| --- | --- | --- | --- | --- |
| Hand | 102.8(3.2) | 4.93(4.3) | 104.6(7.7) | 0.40(2.2) |
| Aim | 97.2(4.5)<sup>n.s.</sup> | -1.42(2.2)<sup>n.s.</sup> | 96.3(6.0)<sup>n.s.</sup> | 0.94(1.8)<sup>n.s.</sup> |
| Implicit | 5.6(6.6)* | 6.35(5.3)* | 8.3(6.0)* | -0.54(2.8)<sup>n.s.</sup> |

The values of aiming locations are compared with the perfect solutions (99.0 mm for x; and 0 mm for y) by simple t tests. The values of implicit adaptations are compared with 0; \*: p<0.05; n.s., p>0.05.

Importantly, the Polar-Based-Correction Model and the Cartesian-Based-Correction Model predicted different time courses of learning for explicit and implicit processes in the perturbation phase. In the Polar-Based-Correction Model, implicit adaptation gradually increased in both x- and y-dimensions (Figure 4A) and reached a considerable amplitude at the end of the perturbation phase. However, the Cartesian-Based-Correction Model predicted that implicit adaptation would only appear in x-dimension, while adaptation in y-dimension would remain zero (Figure 4B). Interestingly, we found implicit adaptation in the rotation and mirror tasks are different. Implicit adaptation in the rotation task were aligned to the prediction of the Polar-Based-Correction Model (Figure 4C). Participants showed considerable implicit adaptation in both x and y-dimensions (X: 5.57 ± 6.61 mm, t_11_ = 2.92, p=0.014; Y: 6.35 ± 5.26 mm, t_11_ = 4.18, p=0.002) under the rotation. On the contrast, implicit adaptation in the mirror task was similar to the prediction of the Cartesian-Based-Correction Model (Figure 4D), where adaptation only appeared in x-dimension (8.27 ± 5.96 mm, t_11_ = 4.81, p=0.0005). A slight negative implicit adaptation was found in y-dimension at the beginning of the mirror task, but it dropped back to zero in the late perturbation (−0.54 ± 2.77 mm, t_11_ = −0.67, p=0.515). Similar to Experiment 1, this adaptation in the mirror task was in the “wrong” direction for mirror reversal which increased the error instead of compensating for it. When comparing between two tasks, the implicit adaptations in x and y-dimensions were roughly equal in the rotation task (t_11_ = 0.267, p=0.794), while they showed a large difference in the mirror task (t_11_ = 3.95, p=0.002). These results indicated that the implicit learning system applied different coordinate systems under different visual perturbations.

To illustrate the adaptation in a more straight-forward way, we presented the aiming location and hand location in the late training of mirror-reversal and visual-rotation, respectively (Figure 5). We get clearly different patterns between the two tasks. In the rotation task, aiming location and hand location fall on a ring with a radius of target distance (Figure 5A), while in the mirror task, the aim and the hand had a large divergence in the x-dimension but little difference in y-dimension (Figure 5B). Thus, it is clear that the learning in the rotation task is more consistent with a polar system, and learning in the mirror task is more consistent with a Cartesian system. Moreover, in the mirror task, the hand overshoot in the x-dimension relative to the aim, indicating that implicit adaptation to mirror-reversal is still in the inappropriate direction.

**Figure. 5.**
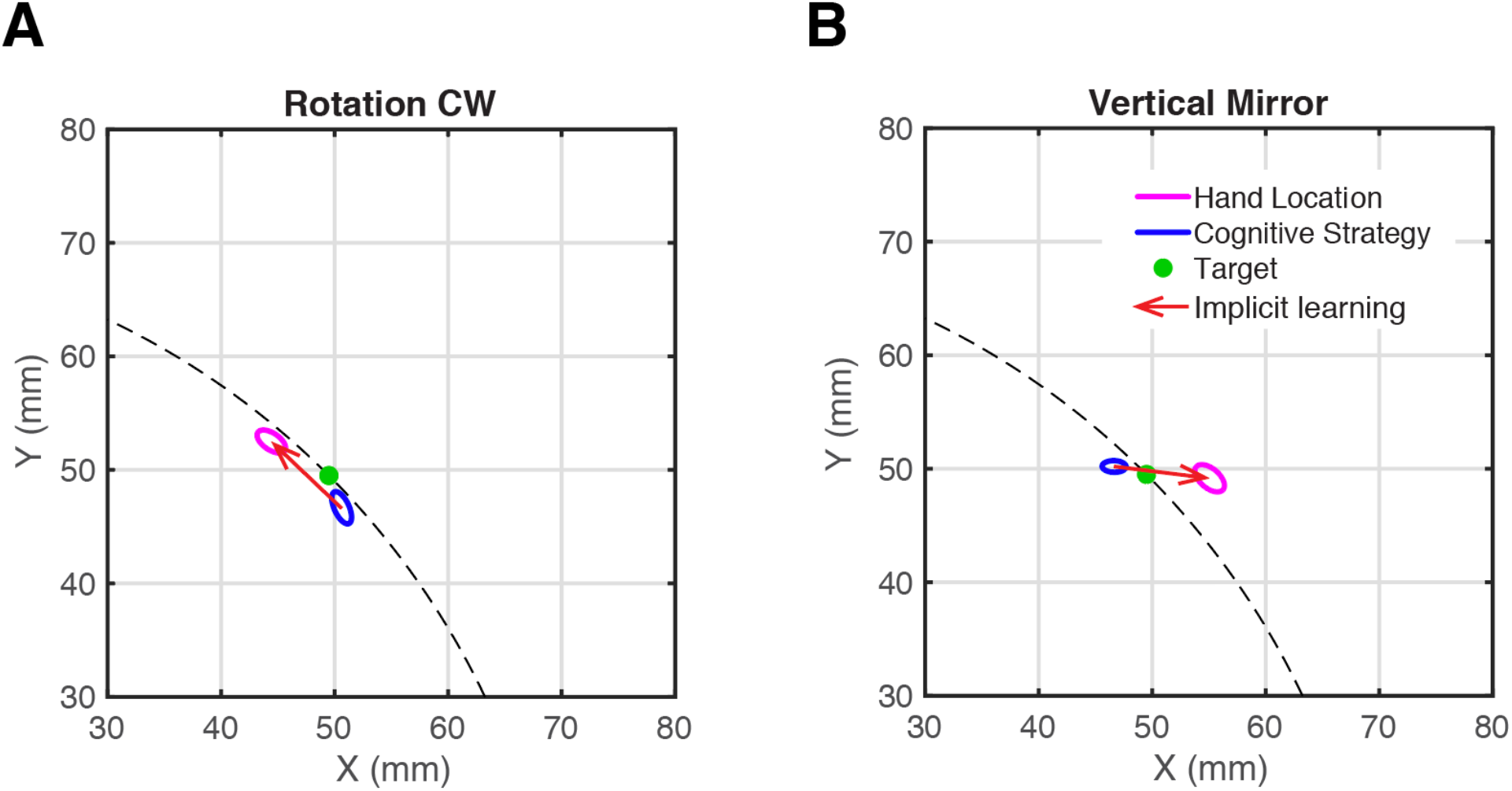
Hand location and aiming location at the end of rotation task and mirror task. A) Covariance ellipses of hand (purple) and aiming location (blue) for 95% CI at the end of the rotation. The data of twelve participants were averaged trial by trial. Hand location and aiming location of all targets are aligned as if the target were in quadrant IV (target −45°, unshown). The covariance of the last forty trials in the perturbation phase was calculated to plot the covariance ellipses. Green spot indicates the target location in quadrant I. B) Covariance ellipses of hand location and aiming location at the end of the vertical mirror. Hand location and aiming location are aligned as if the target were in quadrant II (target 135°, unshown).

### Aftereffect

In both tasks, no significant aftereffect was found in the washout. In rotation task, there was almost nothing left in both x (−0.07 ± 6.42 mm, t_10_ = 0.039, p=0.97) and y-dimension (−2.81 ± 6.80 mm, t_10_ = 1.38, p=0.20) in the washout. In mirror task, hand location slightly went negative in x-dimension (−6.97 ± 7.17 mm, t_10_ = 3.36, p=0.06). Since a correct re-aiming location in training is the same location as another target on the opposite side of the mirror axis, the new aiming location in washout (i.e. directly at the target) is at the same location as that for the opposite target in training. Therefore, the negative aftereffect in x-dimension was aligned with the positive implicit adaptation in the training. Again, very little was found in y-dimension (4.61 ± 5.15 mm, t_10_ = 3.10, p=0.10) in the mirror task. One reason for the weak aftereffect might be we only presented the feedback at the movement end, This kind of feedback was proved to induced less adaptation compared to presenting the feedback during the whole movement (Taylor et al., 2014).

## General Discussion

In the current study, we examined the direction and the coordinate system of implicit adaptation under a mirror reversal perturbation. In the first experiment, we isolated the implicit learning process with clamp mirror perturbation and confirmed that implicit adaptation is in the wrong direction for the mirror reversal, in which the adaptation could only increase the error instead of compensating for it. Interestingly, this adaptation was somehow suppressed during the three-day training. In the second experiment, we examined whether the implicit system was sensitive to the difference in coordinate systems between the rotation and mirror perturbations. This was accomplished by using an aiming report procedure which allowed us to separate implicit and explicit components during the learning. We found the implicit learning system appeared to be consistent with the coordinate system of the particular perturbation, but puzzlingly continued to adapt in the wrong direction for a mirror reversal.

### Adaptation to mirror reversal is in the right coordinate but wrong direction

Both our two experiments aligned with several previous studies that have found that the implicit system learns the mirror reversal in the wrong direction (Gritsenko et al., 2009, Telgen et al., 2014, Kasuga et al., 2015, Hadjiosif et al., 2021, Wilterson & Taylor, 2019). Hadjiosif and colleagues have suggested that this pattern of adaptation in the mirror-reversal is inconsistent with the internal forward model framework for sensorimotor adaptation (Jordan & Rumelhart, 1992; Krakauer & Shadmehr, 2006; Shadmehr, Smith, & Krakauer, 2010). Here, the forward model supposes that the implicit learning system works to minimize the sensory-prediction error (SPE),that is the error between the aim position and the feedback. To reduce an SPE in a mirror reversal, the participant’s hand should adapt closer to the mirror axis, which would bring the cursor and hand to get close to each other. However, the hand appears to adapt in the direction of the rotation and, thus, increases the error in a mirror-reversal task (Wilterson & Taylor, 2019; Hadjiosif et al., 2021). This raised concerns to the internal forward model and asked for other computational models to explain the implicit learning process. Remarkably, even though the implicit component failed to work under mirror reversal, participants actually performed well in the mirror task in Experiment 2 by applying re-aiming strategies. Aligned with previous studies (Wilterson & Taylor, 2019), our result suggested that learning under mirror reversal is at least initially made possible through explicit re-aiming strategies.

Besides the direction of adaptation, the degree of implicit adaptation to different targets in the mirror reversal were consistent with adaptation to different sizes of rotations (Morehead et al., 2017; Schween & Hegele, 2017; Schween et al., 2018). The strong similarity between the performance in mirror and rotation tasks raises the question of whether the implicit system could only process the perturbation as a rotation. However, in a two-dimensional reaching task, in which participants need to control both movement angle and radius, we found implicit adaptation under a mirror reversal was very different from that under a 90° rotation in terms of the coordinate system. This result is consistent with Hudson and Landy’s (2016) study, where they found that participants were sensitive to the coordinate system of an error that was generated in a Polar or a Cartesian system. However, they did not separate the implicit and explicit systems in their study. Our findings suggest that both implicit and explicit learning systems appear to be sensitive to the coordinate system of the perturbation.

We note that, unlike mirror reversal which is somewhat artificial and unlikely to be encountered in everyday life, errors in different coordinate systems may be more common in motor learning. For example, controlling a steering wheel or a bicycle handle asks for manipulation in the Polar system, while a computer mouse may need motor planning in the Cartesian system. However, what information signals the appropriate coordinate system remains to be determined. For every single target in Experiment 2, there was 90° of separation between targets, which is generally outside the range of generalization (Brayanov, Press, & Smith, 2012; Krakauer, Pine, Ghilardi, & Ghez, 2000; Zhou, Fitzgerald, Colucci-Chang, Murthy, & Joiner, 2017). This limitation in the breadth of generalization would suggest that the coordinate system would have to be determined individually at each target location.

There is the possibility that the global pattern of errors could signal the coordinate system of the errors, however, evidence for a meta-learning processes such as this is scant. While incidental contextual cues have been shown to affect learning and change the pattern of generalization (Taylor, Hieber, & Ivry, 2013), it most likely operates on explicit re-aiming. We made sure to remove these visual cues, such as the arrangement of the targets or requiring shooting movements in experiment 2. However, explicit strategies might still operate in different coordinate systems for mirror and for rotation since they ask for different mental manipulations. Different explicit strategies might generate errors in different coordinate systems, which then influences the selection of coordinate system for implicit adaptation. Another possibility is the consistency of the error signals, which has been shown to reduce the sensitivity of implicit adaptation (Castro-Gonzalez, Hadjiosif, Hemphill, and Smith, 2014, Hutter & Taylor, 2018; Albert et al., 2021). In the mirror-reversal task, the sign of the error changes across the workspace while in the rotation task it remains consistent. Again, however, it is unclear if a memory for errors (Herzfeld, Vaswani, Marko, & Shadmehr, 2014) operates globally and could serve as a cue to signal a particular coordinate system.

### Suppression of the implicit adaptation

One interesting result in Experiment 1 is that the adaptation in the wrong direction is suppressed significantly during the three-day training. We also observed suppression of implicit adaptation in our previous five-day contingent mirror task (Wilterson & Taylor, 2019). One explanation for this suppression is that the implicit system changed its error-correction policy during the extended training. When exposed to the mirror reversal, a transformation matrix for rotation was initially applied, which caused the unsuitable adaptation, but after days of training, the implicit system may gradually change the error derivative (Lillicrap et al., 2013; Hadjiosif et al., 2021) or create a more appropriate transformation matrix for a mirror reversal. However, even after days of training, the accumulated adaptation is still in the wrong direction, and to our knowledge, no research has observed an adaptation to the correct direction under mirror reversal. This poses the question of whether the implicit system figure can eventually correct for a mirror reversal or just becomes suppressed.

Attenuation of implicit adaptation does not appear to be specific to the mirror reversal task. Previous studies also found attenuation of the implicit system in some rotation tasks (Avraham et al., 2020). Our previous five-day rotation task (Wilterson & Taylor, 2019) showed a tendency of attenuation in explicit adaptation in the extended training. Moreover, a recent study indicated that implicit adaptation would be attenuated in the re-exposure of the visual perturbation compared to the original adaptation (Leow, Marinovic, de Rugy, & Carroll, 2020; Avraham et al., 2021). What’s more, another study manipulated the variation of the perturbation in a rotation task suggesting that the error sensitivity of the implicit system seems to be suppressed when continuously exposed to the error signal of different directions (Fernandes, Stevenson, & Kording, 2012, Albert et al., 2020). Although the error signal is clamped to each target in our Experiment 1, the error direction is opposite in terms of clockwise or counter-clockwise for targets on the two sides of the mirror axis. Thus, exposure to inconsistent error directions might contribute to the attenuation of implicit adaptation in our result.

### The utility of the implicit system in the sensorimotor learning

The incorrect adaptation direction for mirror reversal and the suppression of the adaptation across days raise the question regarding the utility of implicit adaptation in motor learning. Since the mirror task was indeed the experiment that defined implicit learning (Milner, 1962), it is strange that recent studies showed that implicit adaptation could not overcome the mirror-reversal. A plausible account is that another implicit learning system, e.g., reinforcement learning system, might be critical in motor learning instead of the so-called “implicit adaptation” defined here. The “implicit adaptation” referred to in our analysis was initially defined and proved to be useful in the visuomotor rotation task (Taylor, Krakauer, & Ivry, 2014; Taylor & Ivry, 2011). However, as accumulated evidence showed the limitations of this implicit adaptation (Bond & Taylor, 2015; Hutter & Taylor, 2018; Kim et al., 2018; Schaefer, Shelly, & Thoroughman, 2012, Shmuelof, Krakauer, & Mazzoni, 2012; Morehead et al., 2017), its utility might be restricted to several specific tasks but unable to full fill complex task demands, e.g., mirror-reversal. In those complex situations, this implicit system might be suppressed, and other implicit learning systems somehow help explore and select a proper policy.

### Conclusions

Our study confirmed that implicit adaptation operates in the wrong direction under mirror reversal perturbation. This adaptation was attenuated in the extended training across three days. Although in the wrong direction, the implicit system does appear sensitive to a particular coordinate system. These findings are consistent with previous reports challenging the flexibility of this implicit adaptation process (Bond & Taylor, 2015; Morehead et al., 2017; Wilterson & Taylor, 2019; Hadjiosif et al., 2021).

## Acknowledgments

We thank Eugene Poh and Sarah A. Wilterson for helpful discussions and Chandra Greenberg for helping with data collection. This work was supported by National Institute of Neurological Disorders and Stroke Grant R01 NS-084948 to J. A. Taylor.

## Data availability

The data that support the findings of this study and code for the task and for data analyses are available from the corresponding author upon request.

